# Virtual-SMLM, a virtual environment for real-time interactive SMLM acquisition

**DOI:** 10.1101/2020.03.05.967893

**Authors:** J. Griffié, T.A. Pham, C. Sieben, R. Lang, V. Cevher, S. Holden, M. Unser, S. Manley, D. Sage

## Abstract

Although single molecule localisation microscopy enables for the visualisation of cells nanoscale organisation, its dissemination remains limited mainly due to the complexity of the associated imaging acquisition, impacting on outputs’ reliability and reproducibility. We propose here the first all-in-one fully virtual environment for SMLM acquisition: **Virtual-SMLM**, including on-the-fly interactivity and real time display. It relies on a novel realistic approach to simulate fluorophores photo-physics based on independent pseudo-continuous emission traces. It also facilitates for user-specific experimental and optical environment design. As such, it constitutes a unique tool for the training of both users and machine learning approaches to automated SMLM, as well as for experimental validation, whilst providing realistic data sets for the development of image reconstruction algorithms and data analysis software.

## Main

Super resolution microscopy encompasses a range of diverse imaging techniques that have in common to circumvent the diffraction limit of light, achieving an imaging resolution under 200nm. Since its development, it has opened new avenues of research, allowing for the first time to study cells at the nanoscale with light microscopy. Single molecule localisation microscopy (SMLM) in particular has sparked a growing interest: whilst relying on conventional optical set-ups for the most part as well as existing staining strategies, it uniquely offers a resolution down to 10-30nm[1–4], enabling for quantitative bio-imaging at the molecular level. It relies on the separation of fluorophores’ emission in time (or blinking) recorded over multiple frames. On a given frame, only a small subset of the overall fluorophores population is emitting. It results in spatially well separated point spread functions (PSFs), allowing for precise localisation. Overall, SMLM has proven to be a very powerful tool to identify and investigate cellular structures[5, 6].

Despite these very promising outcomes, SMLM remains far from being a widely used tool across biological sciences. This comes down to a number of issues users are facing: - the complexity of the acquisition process, - the overall lack of guidelines and/ or automation strategies, - the low accessibility to these set-ups for training and optimisation (in terms of price / time). Indeed, although SMLM methodology mostly relies on conventional microscopes (e.g. wide field, total internal reflection), the acquisition itself is highly complex. It includes many user definable parameters (UDP), which impact considerably on the reliability and quality of the output images in a non-intuitive as well as non-linear manner. For new users, this is a considerable setback as tedious and expensive training is required. For instance, both the activation and excitation laser powers have to be set, whilst managing the camera frame rate and gain. Each one of these parameters plays a key role in the fluorophores’ blinking properties and how efficiently it is recorded.

Artefact and limitations (e.g. multiple blinking, high density, relocation over following frames) are directly resulting from the chosen UDP combination[7]. The controversy surrounding protein clusters at the cell membrane is a probing illustration of the limitations of state-of-the-art approaches to SMLM [8–11]. Further complicating, how to best set these UDP depends vastly on the sample itself (e.g. dimensionality, density of the labelling)[12] and remains, for the most part, an open question in the field. As a result, a new sample systematically calls for extensive optimisation and fine tuning on the microscope even for expert users, with no guarantee of results. Overall, these constraints constitute a major flaw when it comes to the reproducibility and reliability of data sets acquired with SMLM, as well as its broader deployment in core biology laboratories. It highlights the crucial need for a realistic and versatile simulation platform allowing, for the first time, to experience a fully virtual SMLM acquisition truly tailored to users’ specific sample and optical set-up, very much like they would experience on a physical microscope. Such platform would not only facilitate users training but also experimental validation and optimisation based on ground truth. It is a requirement for users to investigate the feasibility of novel experiments. Questions such as -“can a given structure be resolved with the experimental design and optical set-up at disposition?,” could be accurately addressed in realistic imaging conditions for the first time.

A very exciting avenue of research that also calls for the development of such virtual acquisition platform is “Smart Microscopy”[13]. It aims at designing self-driving, sample-driven microscopes with limited human inputs. In the case of SMLM, the transition to automated approaches is a requirement if the field wants to provide trustworthy reproducible data. In this context, machine learning (ML) has proven to be an extremely powerful tool to infer optimal UDP combination on-the-fly and is starting to be an active research topic for microscopy, including automated SMLM. Importantly, ML relies heavily on realistic virtual environment for training (e.g. DeepSTORM[14]), which should include real-time AI-simulator interactions and produce a considerable number of studied cases in a short period of time. Self-driving cars for instance are first trained exclusively in such virtual environment, as it leads to reliable future proof as well as fast and safe deployment. In other words, for ML strategies to be applied to SMLM automation, it needs a new simulation tool that recapitulates real life SMLM acquisition.

Whilst some strategies already exist for simulating SMLM like data sets[15–17], they systematically lack the real-time display and the on-the-fly interactivity required for AI and human training. Although more recent studies have started investigating the potentiality of comparing different imaging modalities[18], they continue to lack the requirements of realistic virtual environments. As such, existing simulators have no, or restricted, compatibility with ML training for smart microscopy. Overall, state-of-the-art simulators rely on unrealistic simplified photo-physics (e.g. frame rate dependant, population average) that do not accurately recapitulate the phenomena collected on a physical microscope. They also lack the versatility required for tailored experimental design as well as accurate PSF or camera noise model, and remain limited for the most part to 2D structures. As a result they are only suitable to a very limited number applications, mainly benchmarking or simplistic experimental validation.

We propose here the first virtual platform for SMLM acquisition dedicated to both AI and users training, called: “**Virtual-SMLM**”. This new strategy uniquely enables for real-time display and on-the-fly interaction with the fluorophores photo-physics properties, very much like a user would experience on a physical microscope. As such it provides a milestone for the development and comparison of “smart SMLM” methods. Considerable advances have also been made to the fluorophores photo-physics model, improving significantly the accuracy of the detected fluorescence at the frame level. In addition, to ensure versatility, **Virtual-SMLM** package already includes an intuitive user interface (Supplementary Figure 1), as well as a number of realistic 2D and 3D structures (Supplementary table 1) and imaging modalities (i.e. allowing to simulate a variety of optical set-ups), and it is compatible with any commercially available or home build imaging camera. It is designed to be a fully open-source and community-oriented tool. User can easily add new structures to the range we provide and exchange on how to best image specific structures.

A high level workflow of **Virtual-SMLM** is summarised in Figure 1.a. The input consists of the ground truth structure, defined as a set of 3D coordinates (a.k.a a point cloud) or a surface/volume from which coordinates can be extracted (*online methods*). We already provide a range of such structures but the platform can easily integrate new ground truth data (extracted from electron microscopy (EM) images for instance). Exactly like when planning for a physical experiment, **Virtual-SMLM** includes a number of choices for experimental design, which is divided in two stages: sample preparation and optical set-up. The added value is to generate a virtual acquisition that fits the specificities each user case. For the sample preparation, we propose a range of staining strategies (e.g. direct labelling, immunostaining) (*online methods*). Detailed characteristics concerning the chosen staining method can also be specified if the user has access to them (e.g. labelling efficiency, size of the used antibodies (AB)), resulting in a realistic synthetic sample to image. For the optical set-up, our approach is similar: although we already provide a range of commercially available cameras and illumination solutions (*online methods*), new specifications can easily be uploaded if required. **Virtual-SMLM**’s versatility is a pre-requisite for the development of both efficient analysis and reconstruction software, as well as general ML-based automation strategies for SMLM, as ML relies on realistic and diverse training sets that should mirror real physical case studies.

**Figure 1:**
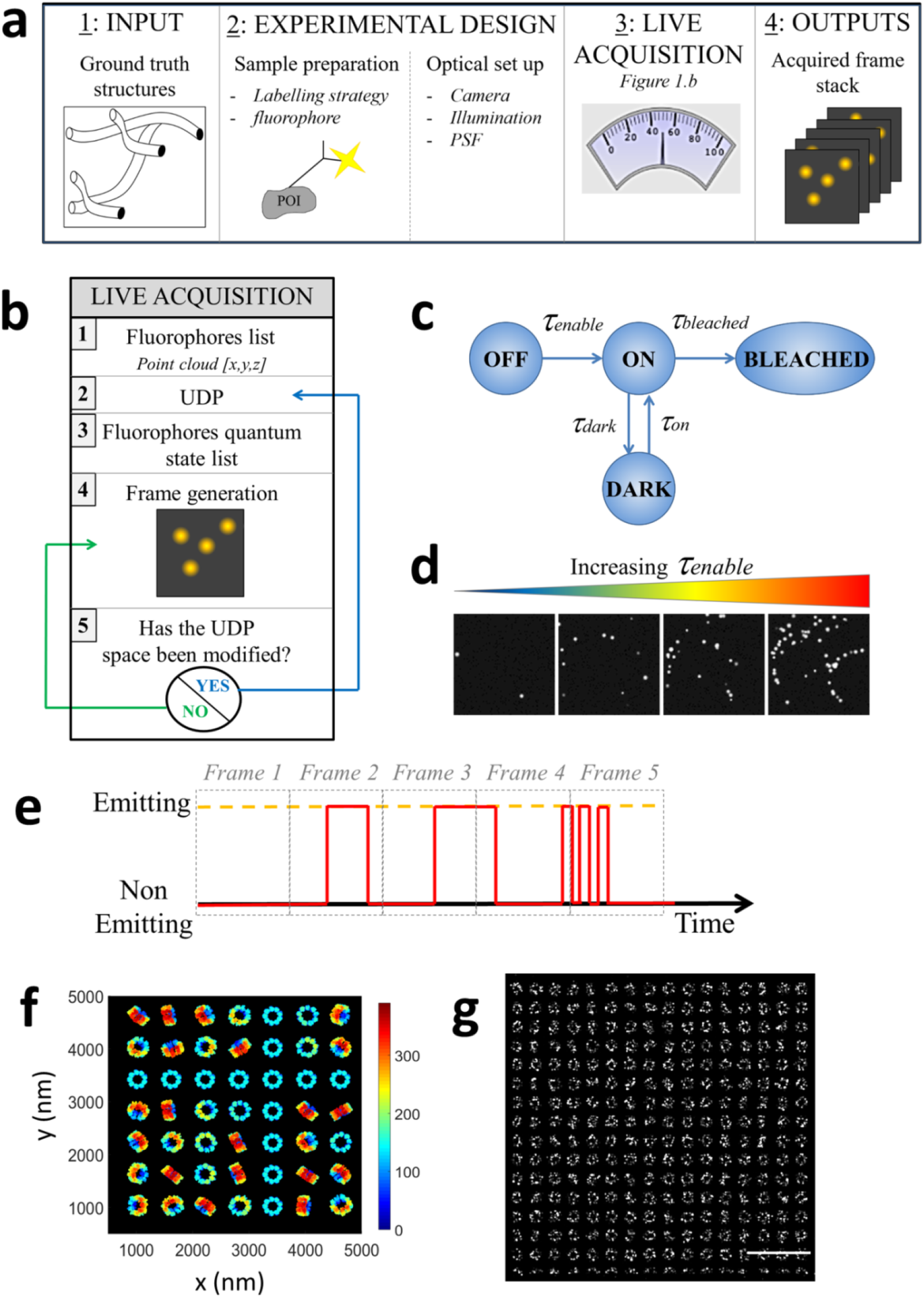
**Virtual-SMLM** methodology. a. high level workflow. POI stands for protein of interest. b. Live acquisition step by step. c. four states model. d. Example frames with fixed proxies except for an increase of τ_enable_. e. Independent pseudo continuous temporal traces for one given fluorophore. f. 2D projection (x,y) of the organization of Cep152 in human centrioles, ground truth point cloud used as input for the virtual imaging session. Colour coded by z point position. g. Resulting super resolved image of the complete FOV following image reconstruction. Cep152 labelled with immunostaining. Scale bar 2μm.

The virtual live acquisition starts once the experimental design is completed. Figure 1.b summarizes the methodology developed for live acquisition. Like on a physical microscope, a number of UDP, which are key to the resulting output image quality, can be modified on-the-fly whilst acquiring. It includes both features associated with camera settings (i.e. frame rate and gain) and laser powers. These parameters are assessed at each ms time step (independently of the frame rate), and uniquely allows for real-time interaction on the virtual microscope. A user can modify the rate of activation for instance, switching from sparsely to densely PSF populated frames on-the-fly. This enables for fine tuning, exactly as would be the case on a physical microscope, whilst providing a unique tool to assess the quality of the reconstructed image versus the ground truth. In parallel, our method ensures real-time display, a requirement for users to assess on-the-fly if the blinking is well recorded and if the PSFs are well enough separated in space to allow for reliable reconstruction. The faithful display required the real-time computation (up to 100Hz) of the convolution with the 2D/3D PSF and the generation of the noise camera.

More specifically, we use a set of proxies to account for the laser powers in our **Virtual-SMLM** microscope. These proxies recapitulate fluorophores photo-physics under given activation and excitation constraints, and are defined by the four states model described in Figure 1c. This model fits accurately photo-activation (PALM) as well as photo-switching (STORM) modes. It relies on a set of 4 rates: τ_enable_, τ_on_, τ_dark_, τ_bleach_, our proxies on the virtual microscope. Concretely, τ_enable_, and τ_on_ are associated with the activation laser, whilst τ_bleach_ and τ_dark_ are linked to the emission laser. We provide examples of frames acquired on **Virtual-SMLM** whilst varying values for the previously described proxies in Figure 1d, and further quantification in Supplementary Figure 2. Although the exact mathematical relationship between these rates and laser powers remains an active topic of research, it is possible to empirically calibrate this relationship for a given experimental design. However, the advantages of using the rate proxies as UDP rather than approximated laser powers are multiple: -it allows for a general framework that do not require for calibration, -it does not impact on ML training, - it facilitates user’s understanding of what they are effectively attempting to fine tuning with the lasers on a physical microscope.

**Virtual-SMLM** also relies on the real-time assessment of each fluorophore’s quantum state. A unique pseudo continuous temporal trace of photo-activity is generated for each fluorophore independently of the frame rate, with a resolution of 1ms (roughly an order of magnitude smaller than the frame rate used for STORM). This is considerable step forward compared to existing simulators which rely on per frame approximation of fluorophores photo-physics (leading to inconsistency in the photon counts and measured uncertainty, as a function of the frame rate chosen), as well as population averaging. In contrast, temporal traces allow for micro-blinks within a frame and spreading of a given fluorophore’s emission over subsequent frames (Figure 1.e). Also, **Virtual-SMLM** relies on Bernoulli statistics for the transition on/dark, dark/on and photo-bleaching, independently for each fluorophore, mimicking accurately a potential change of quantum state at each time step to reach a more realistic scenario (*online methods*). This has been made possible with extensive algorithmic innovation to allow for a significant decrease of associated computer time. We provide the super resolved image reconstructed from a typical virtual imaging session of one of the provided structures, centrioles, using **Virtual-SMLM**(Figure 1.f,g, Supplementary Figure 3 and *online methods*).

To further illustrate the need for virtual environments such as **Virtual-SMLM** to provide reliable and reproducible data, we asked two experts to image 2 consecutive times the same distribution of proteins in PALM mode i.e. synthetic sample and virtual acquisition (*online methods*). We generated for that purpose a ground truth distribution of membrane proteins, with an un-clustered and clustered pool (Figure 2.a), from which we build a realistic synthetic sample using our platform (i.e. immuno-staining). Indeed, although SMLM resolution enables for the visualisation of signalling protein nanoclusters at the cell membrane, it consists of a particularly difficult sample to image. Depending on UDP used for the acquisition, the resulted super resolved image is prone to artefact and lack of reproducibility, giving rise to persistent controversies when used for quantification as previously mentioned.

**Figure 2:**
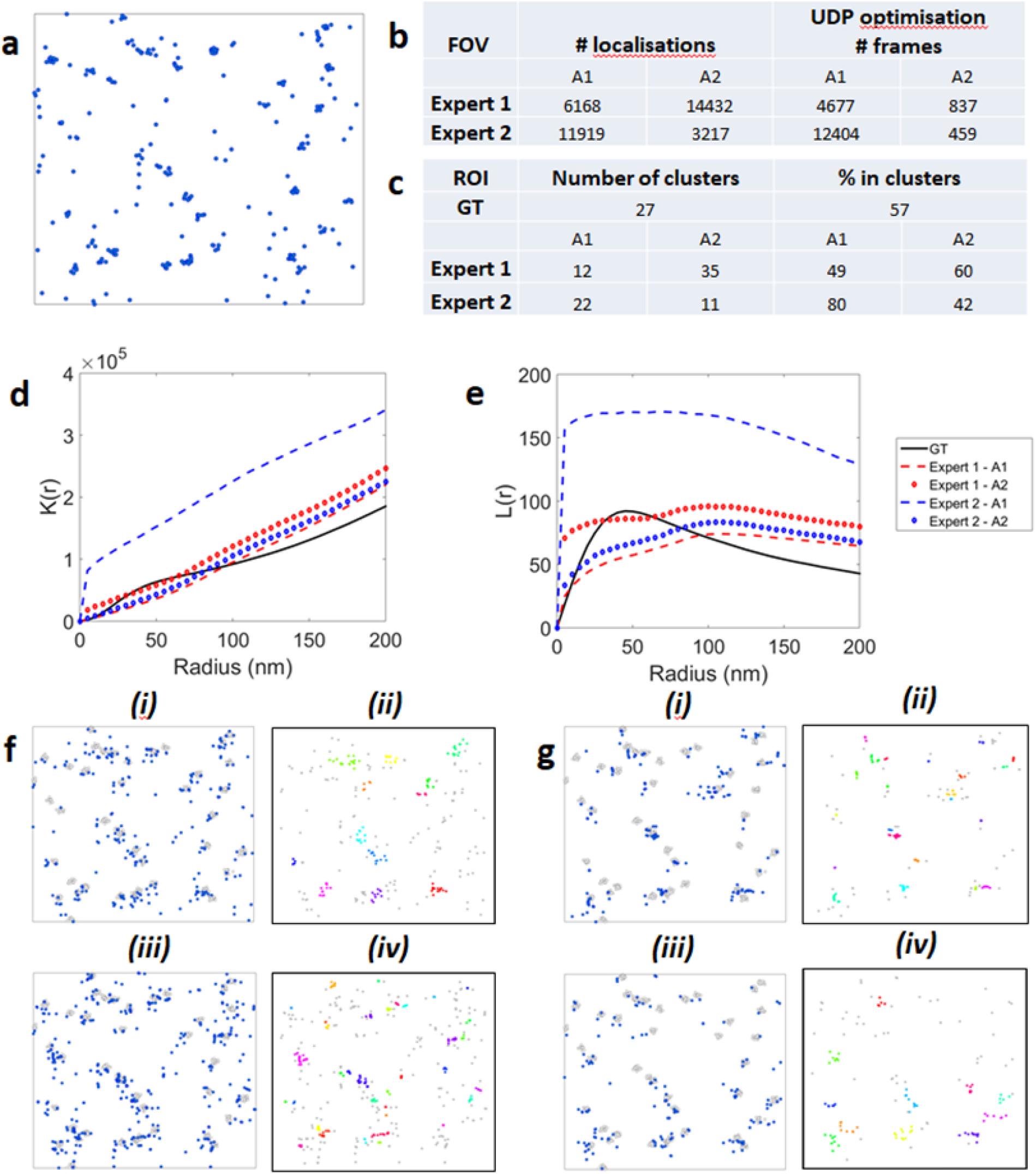
Non-guided virtual SMLM acquisition with two experts. a. Ground truth distribution of proteins for a 2 μm × 2 μm ROI extracted from the FOV. The number of protein has been thinned to mirror the mean number of points collected by for users for the following analysis. b. Overall results (i.e. over the whole FOV) for both experts and both successive acquisitions (A1 and A2). Total number of localisations and time required for fine-tuning the UDP (in number of frames). c. Clustering characteristics extracted from the ROI (e.g. for the ground truth, 57% of proteins were found in clusters, the remaining proteins consists of monomers). d and e Statistical estimation over the whole FOV. d. Ripley’s K estimated curves. e. Linearised Ripley’s L(r) estimated curves. f. 2 μm × 2 μm ROI extracted from the FOV (i) Reconstructed image from Expert 1’s first acquisition in blue, grey for the ground truth clustered proteins and associated (ii) Cluster map (identified clusters are colored, monomers are grey). (iii) Reconstructed image from Expert 1’s second acquisition in blue, grey for the ground truth clustered proteins, and associated (iv) Cluster map. g. (i) Reconstructed image from Expert 2’s first acquisition in blue, grey for the ground truth clustered proteins, and associated (ii) Cluster map. (iii) Reconstructed image from Expert 2’s second acquisition in blue, grey for the ground truth clustered proteins and associated (iv) Cluster map.

Both experts imaged the generated synthetic sample in the same conditions, with 1000 frames recorded for each virtual acquisition. Not only do we observe that the reconstructed localisation distributions vary from one acquisition to the next as the expert optimised or fine-tuned the UDP, but more strikingly, the image quantification for our two experts are drastically different (Figure 2.b-g and Supplementary Figure 4-5). The clustering analysis was extracted using Ripley’s K density curves for entire field of view (FOV) characterisation (Figure 2.d-e) and region of interest (ROI) cluster identification with Bayesian-based cluster analysis tools[8] (Figure 2.c,f,g and *online methods*). We further verified in Supplementary Figure 6 that the observed variations in the reconstructed images, in particular their clustering characteristics, are not resulting from stochastic sampling alone, but rather significant differences on the imaging performances.

The results summarized in Figure 2 are highlighting the limitations of state-of-the-art unguided SMLM. It showcases the need for, at least the rise of consensus and agreed guidelines for SMLM acquisition, and soon user-free ML approaches to super resolution, all of which require virtual environment such as **Virtual-SMLM**. Open source community-driven platform, such as proposed here, directly leads to the rise of novel consensus and guidelines for SMLM as it enables for the development of systematic and robust approaches when it comes to acquisition. With **Virtual-SMLM**, experts can quantitatively compare their methodologies to imaging the same synthetic samples for the first time. Furthermore **Virtual-SMLM** is a requirement for the development of automation strategies relying on benchmark diverse and realistic learning datasets, a necessary transition if SMLM is to increase its dissemination in core biology laboratories. Finally, our approach is fully compatible with multicolour imaging.

## Authors’ contributions

JG and DS designed the project and experiments. JG and DS wrote the manuscript. JG, DS, TAP and RL implemented the software. CS provided biological structures. JG, DS and CS performed simulations. JG, DS SH, SM, MU, VC supervised the project.

## Acknowledgments

Authors acknowledge the following funding sources:

- Zeiss research ideas initiative with EPFL to JG.
- European Research Council (ERC) under European Union’s Horizon 2020 research and innovation program. Grant agreement number 692726 to DS, TAP and MU.
- National Centres of Competence in Research (NCCR) to CS.
- Wellcome Trust & Royal Society Sir Henry Dale Fellowship. Grant number 206670/Z/17/Z to SH.

## Online methods

### Ground truth point clouds from known structures (Supplementary Table 1)

#### Generated via the platform

##### Random distribution (3D)

Proteins of interest are distributed as a complete spatial randomness (CSR) in a volume. The user can provide the number of proteins to throw as well as the dimension (x,y,z) of the volume. This is a particularly useful synthetic sample for 3D imaging modalities.

##### Protein aggregates in a lipid bilayer (2D)

We generate two pools for the protein of interest: clustered and un-clustered proteins in a cellular membrane bilayer. Proteins are projected on a fixed z plane. The clusters’ size, number of enclosed proteins as well as the total number of clusters and percentage of un-clustered proteins are user defined.

#### Provided in the structure package

##### Microtubules (3D)[1]

Microtubules are generated using tubular shapes in the 3D space. These structures are defined by their central axis having typically an average outer diameter of 25 nm with an inner hollow tube of 15 nm diameter. The centre lines of the microtubules are represented by a continuous-domain 3D curve by means of a polynomial spline.

##### Vesicles in a 3D volume[2]

Vesicles are generated as hard edge sphere randomly distributed in a volume. Monomers are overlaid as a CSR in the pre-defined volume. The dimension of the volume, the vesicles’ size, the number of enclosed proteins as well as the total number of vesicles and percentage of un-clustered proteins are user defined. We provide a number of examples in the structures folder.

##### Mixed model: membrane proximal signalling proteins[2]

Signalling proteins proximal to the plasma membrane consist of a mixture model. Both vesicles in the cytosol as well as membrane-bound protein clusters are likely to be present, often associated with a population of monomers in both cases. We provide an example in the structures folder.

##### Centrioles (3D)[3]

The organization of the human centrioles protein Cep152 was modelled into a previously published SMLM 3D reconstruction[3]. To this end, the estimated number of about 550 Cep152 copies[4] was distributed along their 9-fold symmetry into the 3D volume (Supplementary Figure 7).

##### TOROIDs

TOROID ground-truth structures were generated as described previously[5]. Briefly, as a starting model, we generated a double-helix structure with dimensions according to the electron microscopy TOROID reconstruction. The starting structure was then randomly rotated and projected onto the XY plane (Supplementary Figure 7).

### Building a synthetic sample

#### Direct labelling

This scenario models fluorescent proteins. The protein of interest positions are used as fluorophores positions (*in situ*) with a user defined steric hindrance in **Virtual-SMLM** platform.

#### Immuno-staining strategies

Both primary and secondary antibodies (AB) can be used for the immunostaining. The size of the AB, the labelling efficiency as well as the steric hindrance are user defined in **Virtual-SMLM** platform.

For each successive staining step, we generate a new list from our initial list of locations. Each location is moved by the distance of the AB size in a random direction. The probability of a location to be stained (i.e. reported to the newly generated list) is also taken into account at that step (e.g. a labelling efficiency of 0.5 will result in 50% of the locations from the list to not be reported in the new list, a labelling efficiency >1 will result in some locations stained twice). The workflow is summarized in Supplementary Figure 8.

### Optical set up

#### Camera settings

Camera specifications extracted from commercial tools or home-build system can be added manually to the **Virtual-SMLM** platform. The key parameters are: the maximum framerate, the FOV dimension and pixel size, the quantum efficiency, the readout noise and the electron per ADU, as well as the EM gain in the case of an EMCCD camera. All these features are easily accessible on commercial set up user manual.

#### Illumination

We enable for both Gaussian and flat illumination of the FOV, following the development of high-throughput flat-field illumination large FOV[6].

#### PSF

We have used 3D PSFs to realistically generate the 2D shape of the emitter (3D position) in order to compute the 2D frame. These PSF were the ones used in [1]. They were derived from experimentally measured PSFs. for each tested 3D modality: astigmatism, double-helix, biplane and 2D. In practice, the PSF is typically sampled at 25 nm in the three directions and interpolated to the exact positions of the fluorophores.

### Photo physics

For each fluorophore in our synthetic sample, we generate an independent pseudo-continuous temporal trace of emission with 1ms resolution (an order of magnitude smaller than the maximum frame rate used for SMLM). Importantly, these traces are fully independent of the frame rate. We rely on a four states model for the following statistics (Figure 1.c). At each 1ms time point, a small subset of fluorophores is switched from the OFF state to the ON state relying on a binomial distribution. when N (the number of fluorophores) is large and p (probability to turn OFF to ON) is low. Using a binomial law allows to efficiently reproduce the equivalent of a large number of Bernouilli tests for each individual fluorophore. Whilst switched ON a fluorophore is included in the “active fluorophore list”. For every fluorophore entering the active list, we estimate when is the next state transition (from the ON state to either DARK or BLEACHED). For that purpose, we rely on Bernoulli statistics, mimicking quantum state transition mechanism. Which transition a given fluorophore undergoes is set by the chronology of events (the first to transition to test positive on the ms resolved clock). All transitions are independent of the frame rate and can be at any time on the 1ms clock (enabling for micro-blinks). A similar approach is followed for the DARK to ON transition. Once BLEACHED, fluorophores are discarded from the “active” list.

### Application with experts

#### Defining an SMLM expert

We define as an expert in the field of SMLM, a scientist who has had the opportunity to use one or more SMLM set up, on a very regular basis and has had already publications for which SMLM was the main imaging technique used.

#### Imaging

The two experts were presented with the same synthetic sample. Following a tutorial on **Virtual-SMLM** basics, they were asked to image the synthetic sample twice in the same conditions than they would have experienced on a physical microscope.

The ground truth (GT) protein distribution is provided in the Structures folder. It consists of a distribution of proteins in a membrane bilayer (z=0), including clusters of proteins and monomers. To switch from GT to synthetic sample, we designed a 2 stages immuno-staining directly on **Virtual-SMLM** platform:

- primary AB of 15nm with a labelling efficiency of 90%
- secondary AB of 15nm with a labelling efficiency of 100%

For the Optical set up, we used the specifications of the Evolve EMCCD 100×100 Delta and a Gaussian illumination of the sample. We performed a conventional 2D imaging virtual session. For each acquisition 1000 frames of 30ms were recorded.

#### Overall quantification

For the overall quantification of the experts’ imaging performances, we relied on ThunderSTORM[7] to extract the fluorophores localisations from the collected frames. Multi emitter fitting was performed to facilitate the detection of overlapping point spread functions, a very likely phenomenon in the context of a clustered distribution. We then estimated the total number of localisations extracted from each acquisition, the time the experts required for the optimisation/fine-tuning of the acquisition related user defined parameters (UDP) as well as the final choice of UDP for each acquisition. Furthermore, overall clustering characteristics were estimated relying on Ripley’s K curves.

#### Cluster analysis

A region of interest (ROI) of 2μm × 2 μm was extracted from the field of view for both experts’ acquisitions and the GT distribution. The GT ROI was thinned down to a comparable number of points than collected by the experts to allow for fair comparison. We performed state-of-the-art Bayesian based cluster analysis relying on topographic prominence for its thresholding[2,8]. The results are striking, in particular the impact of the sample preparation (i.e. the use of AB) on the cluster size as well as the dramatic effect of UDP on the final cluster maps. From these cluster maps a number of parameters can be extracted: the number of clusters, their size and the number of proteins enclosed per clusters as well as overall characteristics such as the percentage of proteins in clusters and the number of clusters per ROI.

## References

1. Betzig, E., et al., Imaging Intracellular Fluorescent Proteins at Nanometer Resolution. Science, 2006. 313(5793): p. 1642–1645.

2. Hess, S.T., T.P.K. Girirajan, and M.D. Mason, Ultra-high resolution imaging by fluorescence photoactivation localization microscopy. Biophysical journal, 2006. 91(11): p. 4258–4272.

3. Rust, M.J., M. Bates, and X. Zhuang, Sub-diffraction-limit imaging by stochastic optical reconstruction microscopy (STORM). Nat Methods, 2006. 3(10): p. 793–5.

4. Jungmann, R., et al., Single-Molecule Kinetics and Super-Resolution Microscopy by Fluorescence Imaging of Transient Binding on DNA Origami. Nano Letters, 2010. 10(11): p. 4756–4761.

5. Sieben, C., et al., Multicolor single-particle reconstruction of protein complexes. Nature Methods, 2018. 15(10): p. 777–780.

6. Holden, S.J., et al., High throughput 3D super-resolution microscopy reveals Caulobacter crescentus in vivo Z-ring organization. Proceedings of the National Academy of Sciences, 2014. 111(12): p. 4566–4571.

7. Burgert, A., et al., Artifacts in single-molecule localization microscopy. Histochemistry and Cell Biology, 2015. 144(2): p. 123–131.

8. Griffié, J., et al., A Bayesian cluster analysis method for single-molecule localization microscopy data. Nature Protocols, 2016. 11(12): p. 2499–2514.

9. Levet, F., et al., SR-Tesseler: a method to segment and quantify localization-based super-resolution microscopy data. Nature Methods, 2015. 12(11): p. 1065–1071.

10. Rossboth, B., et al., TCRs are randomly distributed on the plasma membrane of resting antigen-experienced T cells. Nature Immunology, 2018. 19(8): p. 821–827.

11. Griffié, J., et al., Dynamic Bayesian Cluster Analysis of Live-Cell Single Molecule Localization Microscopy Datasets. Small Methods, 2018. 2(9): p. 1800008.

12. Fox-Roberts, P., et al., Local dimensionality determines imaging speed in localization microscopy. Nature communications, 2017. 8: p. 13558–13558.

13. Scherf, N. and J. Huisken, The smart and gentle microscope. Nature Biotechnology, 2015. 33(8): p. 815–818.

14. Nehme, E., et al., Deep-STORM: super-resolution single-molecule microscopy by deep learning. Optica, 2018. 5(4): p. 458–464.

15. Sage, D., et al., Super-resolution fight club: assessment of 2D and 3D single-molecule localization microscopy software. Nature Methods, 2019. 16(5): p. 387–395.

16. Venkataramani, V., et al., SuReSim: simulating localization microscopy experiments from ground truth models. Nature Methods, 2016. 13(4): p. 319–321.

17. Novák, T., et al., TestSTORM: Versatile simulator software for multimodal super-resolution localization fluorescence microscopy. Scientific reports, 2017. 7(1): p. 951–951.

18. Lagardère, M., et al., FluoSim: simulator of single molecule dynamics for fluorescence live-cell and super-resolution imaging of membrane proteins. bioRxiv, 2020: p. 2020.02.06.937045.

## References

1. Sage D, Pham T-A, Babcock H, Lukes T, Pengo T, Chao J, et al. Super-resolution fight club: assessment of 2D and 3D single-molecule localization microscopy software. Nature methods. 2019;16(5):387–95. doi: 10.1038/s41592-019-0364-4.

2. Griffié J, Shlomovich L, Williamson DJ, Shannon M, Aaron J, Khuon S, et al. 3D Bayesian cluster analysis of super-resolution data reveals LAT recruitment to the T cell synapse. Sci Rep. 2017;7(1):4077. doi: 10.1038/s41598-017-04450-w.

3. Sieben C, Banterle N, Douglass KM, Gönczy P, Manley S. Multicolor single-particle reconstruction of protein complexes. Nature methods. 2018;15(10):777–80. doi: 10.1038/s41592-018-0140-x.

4. Bauer M, Cubizolles F, Schmidt A, Nigg EA. Quantitative analysis of human centrosome architecture by targeted proteomics and fluorescence imaging. The EMBO journal. 2016;35(19):2152–66. Epub 08/18. doi: 10.15252/embj.201694462. PubMed PMID: 27539480.

5. Prouteau M, Desfosses A, Sieben C, Bourgoint C, Lydia Mozaffari N, Demurtas D, et al. TORC1 organized in inhibited domains (TOROIDs) regulate TORC1 activity. Nature. 2017;550(7675):265–9. doi: 10.1038/nature24021.

6. Douglass KM, Sieben C, Archetti A, Lambert A, Manley S. Super-resolution imaging of multiple cells by optimised flat-field epi-illumination. Nat Photonics. 2016;10(11):705–8. Epub 10/17. doi: 10.1038/nphoton.2016.200. PubMed PMID: 27818707.

7. Ovesný M, Křížek P, Borkovec J, Svindrych Z, Hagen GM. ThunderSTORM: a comprehensive ImageJ plug-in for PALM and STORM data analysis and super-resolution imaging. Bioinformatics. 2014;30(16):2389–90. Epub 04/25. doi: 10.1093/bioinformatics/btu202. PubMed PMID: 24771516.

8. Griffié J, Shannon M, Bromley CL, Boelen L, Burn GL, Williamson DJ, et al. A Bayesian cluster analysis method for single-molecule localization microscopy data. Nature Protocols. 2016;11(12):2499–514. doi: 10.1038/nprot.2016.149.

