## Supplementary Information for "Virtual-SMLM, a virtual environment for real-time interactive SMLM acquisition"

### Supplementary figures

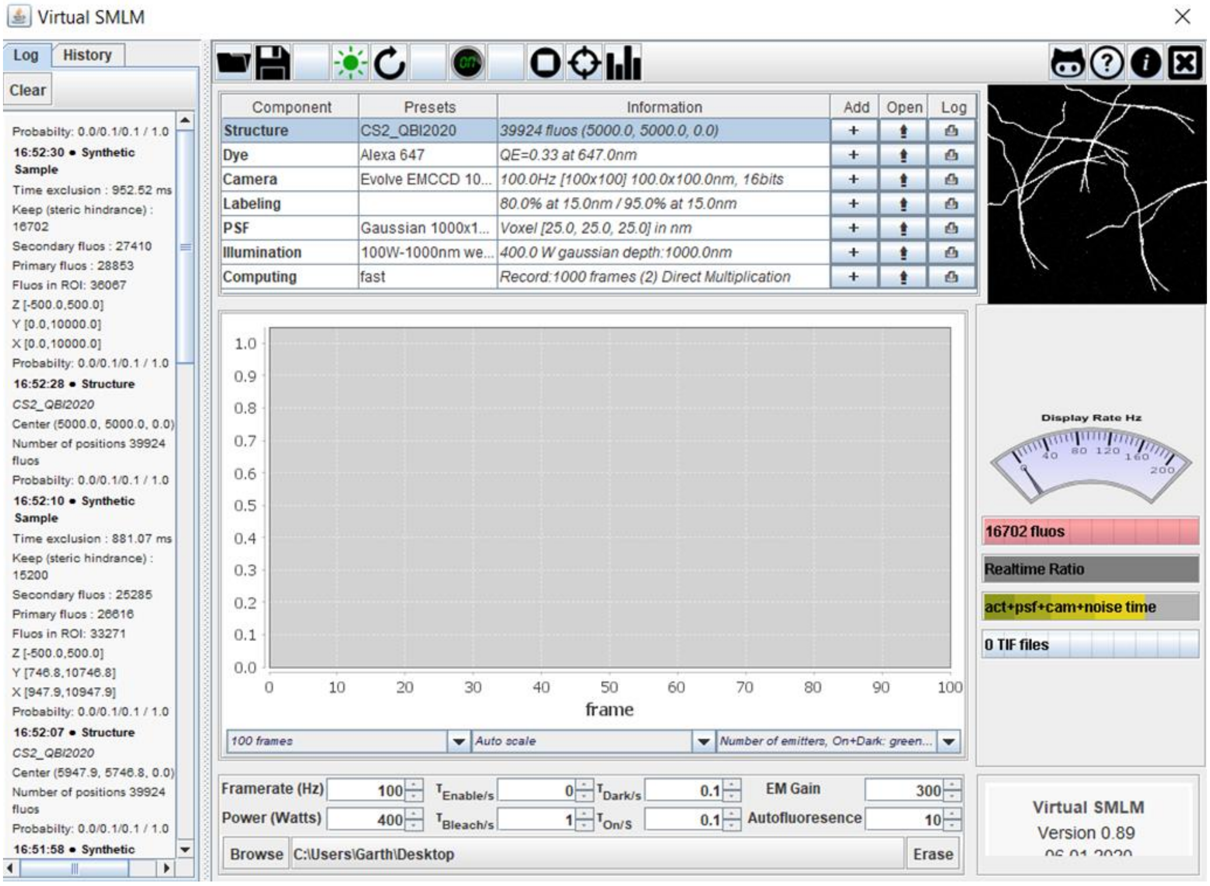


**Supplementary Figure 1: Virtual-SMLM** user interface.

**a**

| | t0 (b) | t1 (c) | t2 (d) | t3 (e) | t4 (f) |
|---|---|---|---|---|---|
| τenable | 20 | 70 | 150 | 150 | 150 |
| τbleach | 20 | 20 | 20 | 70 | 150 |

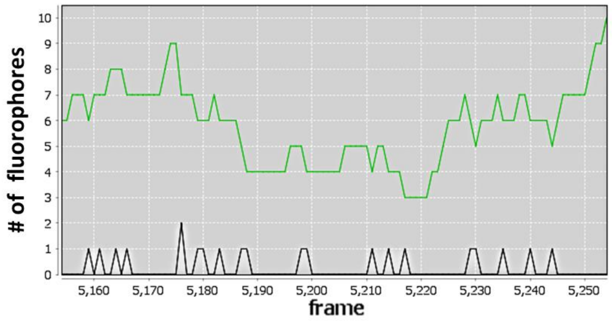


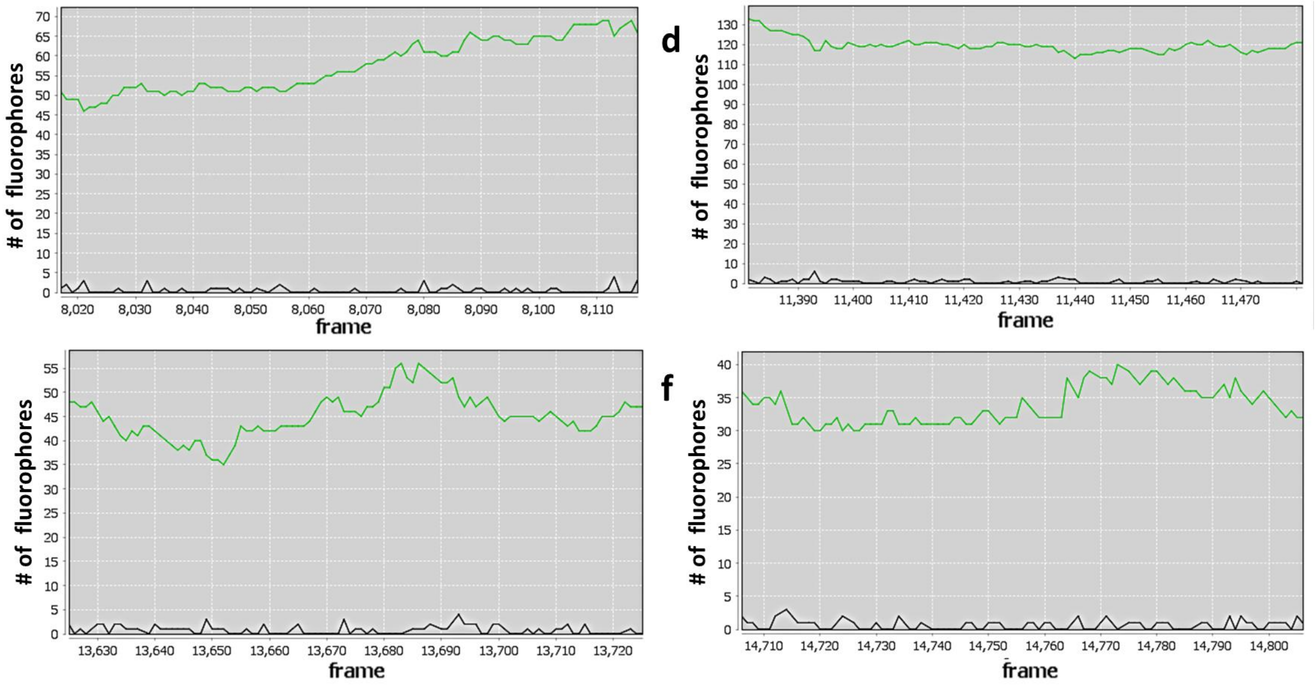


**Supplementary Figure 2:** Illustration of UDP's impact on the number of active i.e. on and dark states combined (in green) and bleached (in black) fluorophores per frame. a. Table recapitulating $\tau_{bleach}$ and $\tau_{enable}$ used for the following measurements. b,c,d,e,f Optimisation curves provided on **Virtual-SMLM** platform. Green: counts for active fluorophores (i.e. in on and dark states combined), grey counts for the bleached fluorophores per frame.

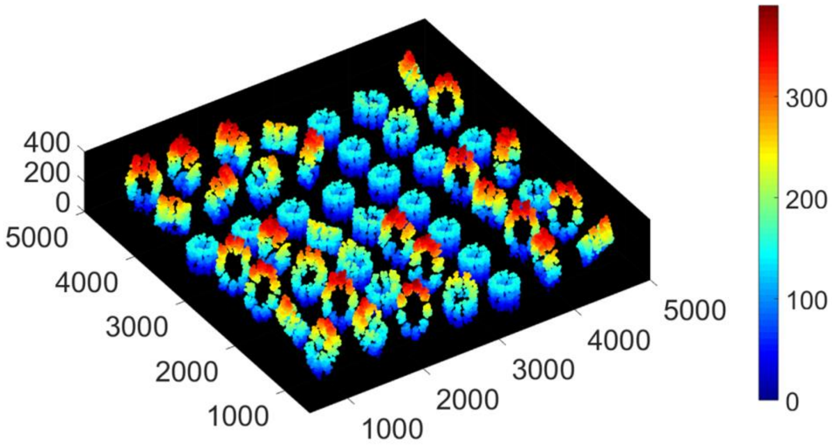


**Supplementary Figure 3:** Ground truth point cloud. 3D organization of Cep152 in human centrioles. Colour coded with z point position. (x,y,z) in nm.

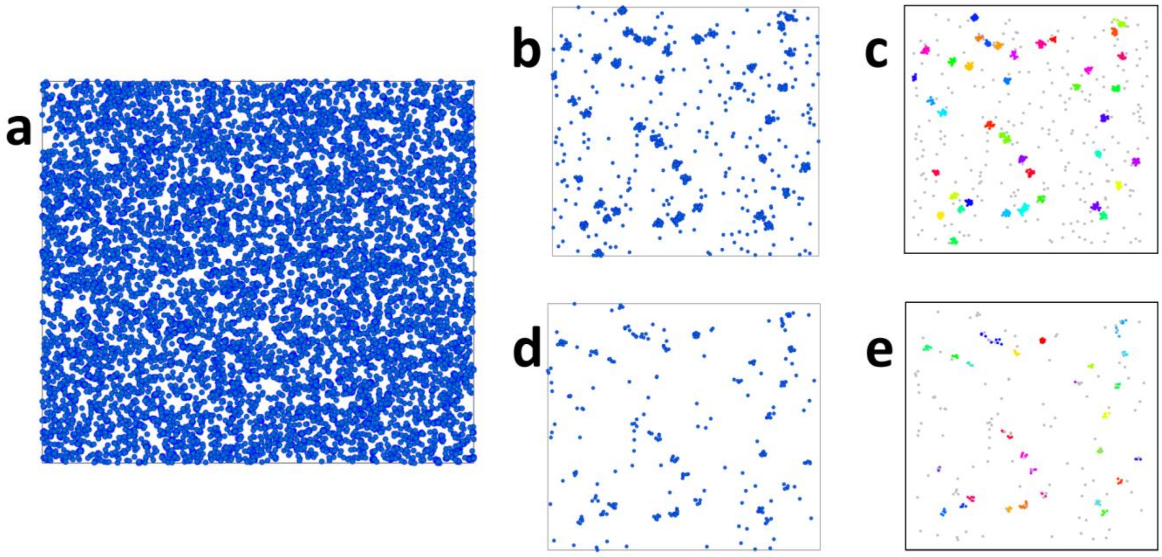


**Supplementary Figure 4:** Ground truth (GT) protein distribution. a. distribution over the whole field of view (FOV). b. 2µm x 2µm region of interest (ROI) extracted from the GT distribution and c. associated cluster map. d. GT ROI with 35 % of proteins (to mimick the collected number of localisations by Expert 1 and 2 over 1000 frames) and e. associated cluster map.

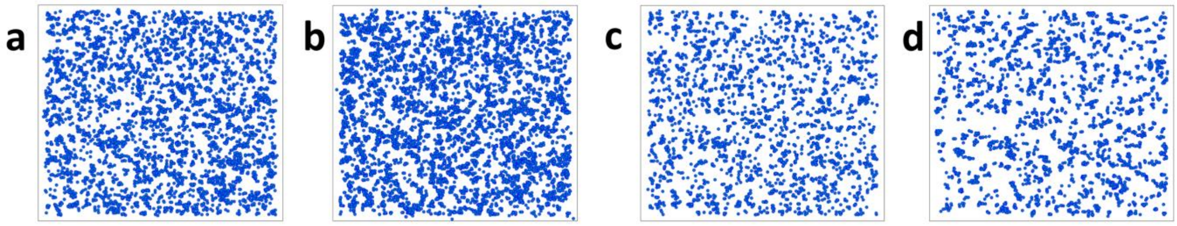

**Supplementary Figure 5:** Results over the whole FOV for expert 1 and 2. a. Expert 1 Acquisition 1: distribution over the whole FOV. b. Expert 1 Acquisition 2: distribution over the whole FOV. c. Expert 2 Acquisition 1: distribution over the whole FOV. d. Expert 2 Acquisition 2: distribution over the whole FOV.

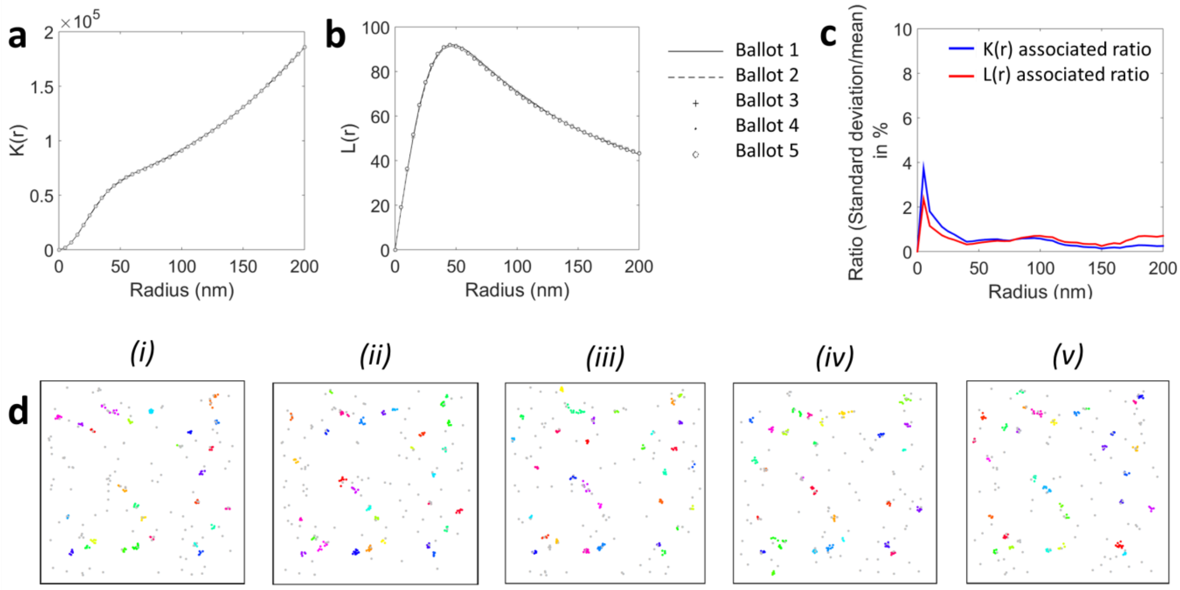


**e**

| ROI | Number of clusters | | % in clusters | |
|---|---|---|---|---|
| n=5 | Mean | σ | Mean | σ |
| | 29.6 | 1.8 | 60.4 | 2.1 |

**Supplementary Figure 6:** Stochastic sampling impact on cluster characteristics. a, b and d concern overall clustering characteristics over the whole FOV. The ground truth distribution was thinned down to a comparable number of points than acquired by the experts, in five independent ballots, illustrating stochastic sampling. a. The Ripley's K curves for the 5 different ballots are nearly perfectly overlaid. b. The Ripley's L curves for the 5 different ballots are nearly perfectly overlaid. c. Ratio of the standard deviation divided by the mean of the calculated K and L curves over the 5 independent ballots. d and e concern full clustering characteristics over a ROI. The ground truth distribution was thinned down to a comparable number of points than acquired by the experts, in 5 independent ballots, illustrating stochastic sampling. d. Resulting cluster maps over the five independent ballots e. Quantification of the clustering characteristics extracted from the cluster maps.

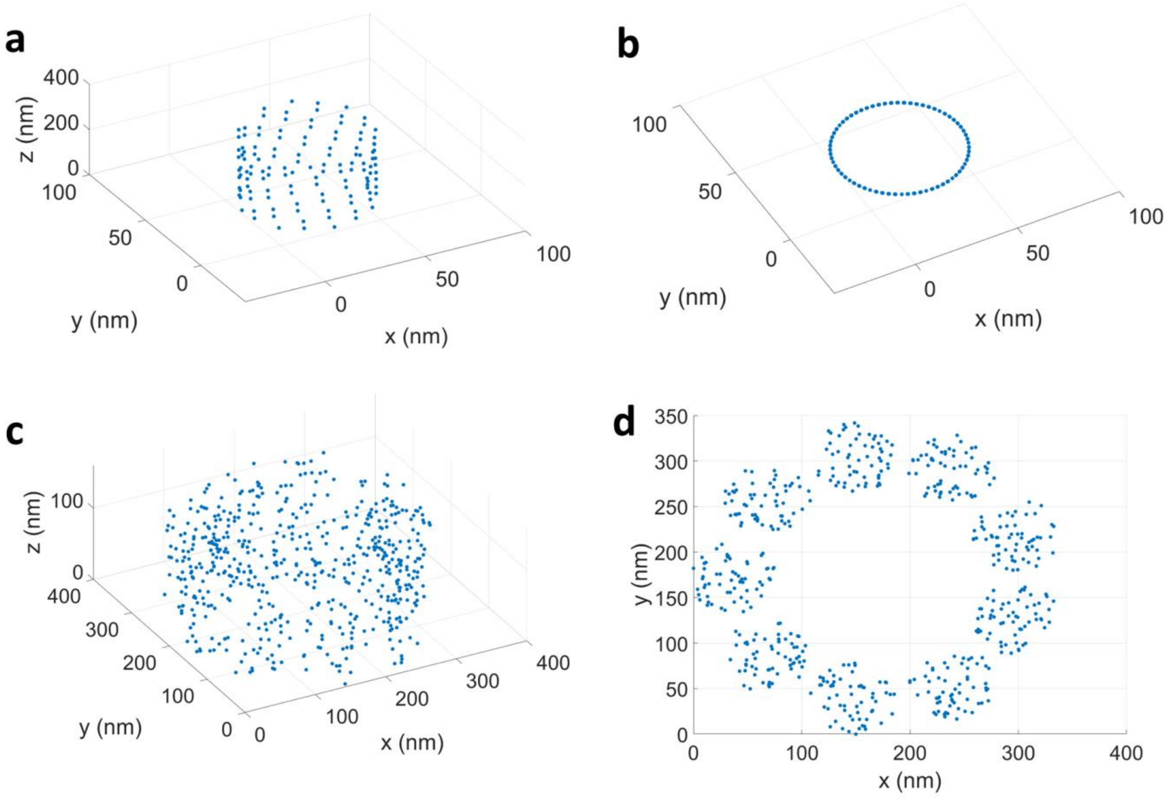


**Supplementary Figure 7:** Example of provided structures extracted from known molecular structures. a. 3D view of a TOROID. b. 2D view of a TOROID. c. 3D view of Cep152 in human centrioles. d. 2D projection (x,y) of the organization of Cep152 in human centrioles.

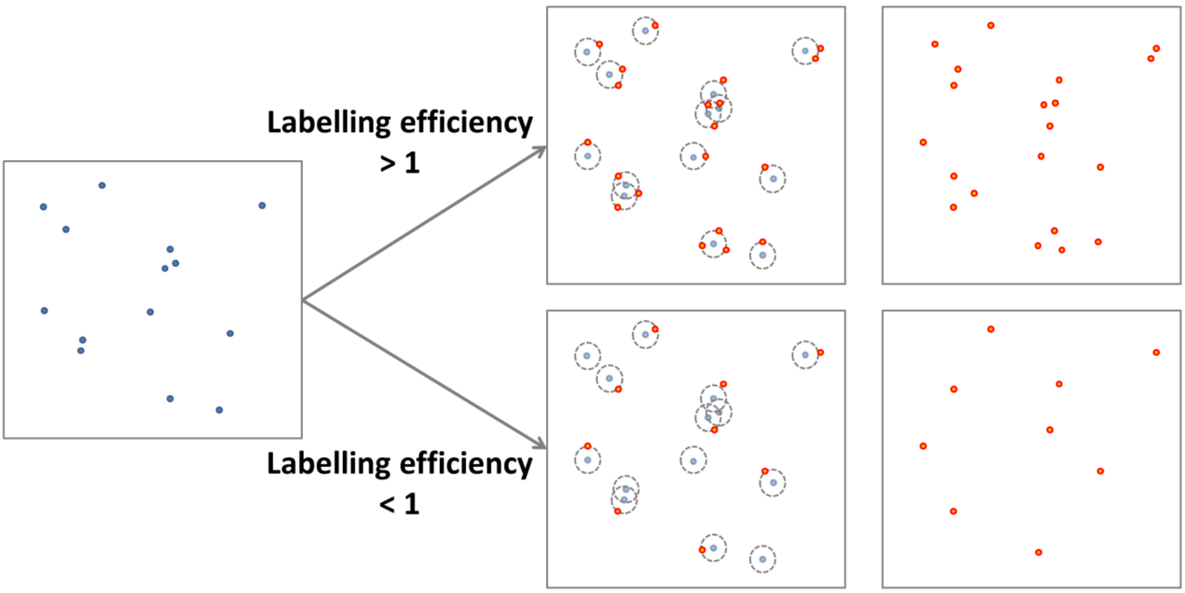


**Supplementary Figure 8:** Experimental design with the implementation of immuno-staining principle. Simplified 2D methodology schematic. In blue the GT distribution of protein. In red is the resulting distribution after the use of primary AB for the targeting of the protein of interest, with 2 different scenarios for the labelling efficiency.

### Supplementary table

| | 2D | 3D | Real data based | Model based | Generated | Provided |
|---|---|---|---|---|---|---|
| CSR in a volume | | X | | X | X | |
| Clusters in membrane | X | | | X | X | |
| Microtubules | | X | X | | | X |
| Vesicles | | X | | X | | X |
| Membrane proximal signalling | | X | | X | | X |
| Centrioles | | X | X | | | X |
| TOROIDs | | X | X | | | X |

**Supplementary Table 1:** Structures included in **Virtual-SMLM** package. Some structures can be generated directly from the platform (CSR in a volume and clusters in a membrane), the rest are provided in the structure package and can be uploaded to the interface as well.

### Code availability

Full source code will be available at a dedicated repository (on corresponding author's laboratory GitHub) under an approved open source license at the time of journal publication.

### Data availability

The ground truth distribution used for the applications with experts as well as the other discussed provided structures can be found in the structure folder of **Virtual-SMLM** package.
